# BRAID: RT-PCR-calibrated conformal intervals for splicing ΔPSI

**DOI:** 10.64898/2026.07.02.736001

**Authors:** Jihye Park, Keunsoo Kang

**Affiliations:** Department of Biomedical Sciences, College of Bio-Convergence, Dankook University, Cheonan 31116, Republic of Korea; MIND-X Superintelligence Convergence Innovation Lab, Dankook University, Cheonan 31116, Republic of Korea

## Abstract

Differential splicing workflows usually report a ΔPSI point estimate and a statistical score, but these outputs do not directly state whether the RNA-seq estimate is close enough to an orthogonal validation measurement. We developed BRAID as a post-processing calibration step for splicing analyses. BRAID estimates RNA-seq ΔPSI from rMATS inclusion and skipping counts, retains the upstream caller evidence, and adds a 95% interval whose width is calibrated from empirical RNA-seq-to-RT-PCR residuals using split conformal prediction. The packaged differential-splicing calibrator uses a residual half-width of *q* = 0.341, estimated from 162 RT-PCR-validated skipped-exon events. We evaluated BRAID on three RT-PCR validation datasets covering *TRA2* knockdown, mouse cerebellum versus liver, and a prostate epithelial-to-mesenchymal comparison. On the pooled common set of 139 cassette-exon events, BRAID reached 0.971 RT-PCR coverage, whereas MAJIQ, betAS, and rMATS-derived intervals reached 0.518, 0.734, and 0.633, respectively. BRAID also had the lowest pooled interval score, 0.720, compared with 2.040 for MAJIQ, 1.414 for betAS, and 1.625 for rMATS. Applying the same residual calibration to other caller outputs brought MAJIQ, betAS, rMATS, and SUPPA2 ΔPSI estimates close to nominal RT-PCR coverage, indicating that the gain came from interval calibration rather than from a caller-specific point estimate. In a *TRA2* positive-negative validation panel, using q as a hard rMATS effect-size cutoff reduced recall, whereas using q as an interval half-width improved RT-PCR coverage. Applied to a public DM1 skeletal-muscle rMATS table, BRAID reduced 967 large-effect significant events to 68 high-confidence interval-supported events and retained known DM1 and muscle-splicing signals. BRAID provides a practical calibrated reliability layer for RNA-seq splicing studies where downstream follow-up depends on the precision of reported ΔPSI estimates.

## Introduction

Alternative splicing expands transcript diversity and shapes gene regulation across development, tissue identity, and disease [1,2]. Short-read RNA-seq has made transcriptome-wide splicing analysis routine, including event-level quantification across human tissues [3]. For cassette-exons, splicing changes are commonly summarized as ΔPSI, the difference in percent-spliced-in (PSI) between two conditions; PSI estimates the fraction of transcripts that include the exon rather than skip it [4]. Tools such as rMATS, MAJIQ, SUPPA2, and betAS estimate differential splicing from RNA-seq data, with outputs that include event-level effect estimates, statistical scores, and, for some methods, RNA-seq-derived uncertainty summaries [4–8]. For experimental follow-up, event ranking is insufficient when downstream decisions depend on whether the RNA-seq ΔPSI estimate is close to an orthogonal validation measurement.

These outputs support detection and ranking, but they do not by themselves answer whether an interval centered on the RNA-seq estimate will cover an orthogonal RT-PCR ΔPSI measurement. This distinction matters because RNA-seq and RT-PCR differ in assay chemistry, primer or junction definition, length normalization, and event modeling. Becuase RNA-seq and RT-PCR quantify related but non-identical molecular summaries, RNA-seq model uncertainty does not necessarily represent agreement with RT-PCR validation measurements [9].

BRAID addresses this gap as a post-processing calibration step, not as a new splicing caller. In the rMATS workflow, BRAID reads existing event tables, keeps the caller-derived ΔPSI estimate, and adds a calibrated interval and confidence tier. The interval half-width is learned from absolute RNA-seq-to-RT-PCR residuals using split conformal prediction [10]. Under exchangeability, this procedure provides distribution-free marginal coverage for future events drawn from the same calibration population, while leaving the upstream splicing workflow unchanged.

Here, we evaluate whether adding a calibrated residual width to existing RNA-seq ΔPSI estimates yields intervals with nominal RT-PCR coverage. We test this across three RT-PCR coverage datasets, using native method intervals when available and intervals reconstructed from posterior summaries when only posterior summaries are reported. This design keeps the caller estimate fixed while shifting the uncertainty target from RNA-seq model fit to RT-PCR coverage.

## Materials and methods

### Study design

BRAID was evaluated on event-level differential alternative-splicing results. The target quantity was ΔPSI, defined as the difference in percent-spliced-in (PSI) between the two conditions after orienting each dataset to the corresponding RT-PCR contrast. The primary endpoint was empirical coverage: the proportion of events for which a nominal 95% interval centered on the RNA-seq ΔPSI estimate contained the paired RT-PCR ΔPSI measurement. Mean interval width and the Gneiting-Raftery interval score [11] were used to summarize interval sharpness.

### rMATS-derived ΔPSI

The rMATS workflow used standard junction-count tables (‘*.MATS.JC.txt’) as input [4]. By default, rMATS sample 1 (‘--b1’) was treated as the first condition and sample 2 (‘--b2’) as the second condition. ΔPSI was therefore defined as sample 1 minus sample 2, matching rMATS IncLevelDifference. When the paired RT-PCR table used the opposite contrast, the sign of ΔPSI was reversed before scoring; the same orientation was then used for all methods.

For each skipped-exon event, let *I* and *S* denote the inclusion-junction and skipping-junction counts, and let *L_I_* and *L_S_* denote the corresponding rMATS inclusion and skipping form lengths. PSI was computed on the rMATS IncLevel scale as:

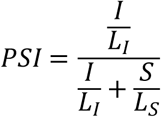

Equivalently, BRAID kept the inclusion count unchanged and rescaled the skipping count as:

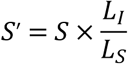

For the pooled model, group-level PSI was sampled from a Jeffreys-Beta posterior:

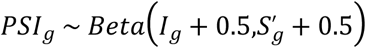

where *g* indexes the condition. ΔPSI samples were then obtained as:

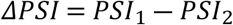

The command-line default used 500 posterior draws. If complete per-replicate rMATS count vectors were available in both groups, the auto model resampled replicate pairs and averaged replicate-level PSI draws; otherwise it used pooled group counts. The main benchmark analyses used the pooled ΔPSI surface.

### RT-PCR-calibrated interval

For calibration event *i*, let *m_i_* denote the RNA-seq ΔPSI estimate and *y_i_* denote the paired RT-PCR ΔPSI measurement. The nonconformity score was defined as the absolute residual:

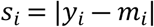

For target miscoverage *α*, the split-conformal half-width *q* was set to the residual with rank equal to the smallest integer greater than or equal to (*n* + 1)(1 − *α*) among *n* calibration events. For a new event with RNA-seq estimate *m*, the 95% interval was [*m* − *q*, *m* + *q*], clipped to the valid ΔPSI range [−1, 1]. If the conformal rank exceeded the calibration-set size, the interval was set to the full valid range. At *α* = 0.05, a finite rank requires at least 19 calibration events.

The primary RT-PCR benchmark used five-fold cross-fit calibration. For each fold, *q* was estimated from the other folds and applied to the held-out fold. Support-stratified Mondrian calibration used the bins <20, 20–49, 50–99, 100–249, and 250+, with a pooled fallback for sparse bins.

The package includes a default differential ΔPSI calibrator fitted on 162 skipped-exon events with RT-PCR truth from the *TRA2* and circadian surfaces. The global half-width is *q* = 0.341 at *α* = 0.05. In the braid differential command, this residual width is combined with RNA-seq sampling spread as:

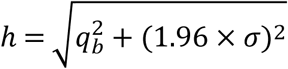

where *q_b_* is the selected conformal quantile and *σ* is the event-level ΔPSI sampling standard deviation.

### Confidence tiers and caller wrappers

BRAID reports the calibrated interval together with upstream caller evidence and a confidence tier. An event was defined as reliable when the calibrated interval excluded zero. The default effect threshold was |ΔPSI| ≥ 0.1, and the default caller-significance threshold was 0.05 for FDR or P value. Events satisfying all three criteria were labeled high-confidence. Reliable effect-size events not significant by the caller were labeled supported, and caller-significant events whose calibrated interval crossed zero were labeled caller-significant-only.

The braid filter command maps rMATS, MAJIQ, SUPPA2, and betAS outputs into a common event record [4–8]. The record includes the event identifier, gene, event type, ΔPSI estimate, caller significance fields, support when available, and native interval bounds when present. Callers without countable support were assigned the pooled global conformal quantile. In caller-specific evaluations, *q* was fitted or cross-fitted on residuals from the corresponding caller and validation surface.

### RT-PCR datasets and event matching

Three RT-PCR validation datasets were used. The *TRA2* dataset used GSE59335, a human MDA-MB-231 *TRA2A/TRA2B* knockdown experiment with three control and three knockdown RNA-seq replicates; the matched rMATS set contained 112 events, and the four-method common set contained 62 [7,12]. The circadian dataset used GSE54651, comparing mouse cerebellum and liver replicates, and contained 50 RT-PCR ΔPSI targets [6,7,13]. The PC3E/GS689 dataset used the human epithelial-to-mesenchymal contrast from the rMATS study and contained 34 RT-PCR-positive events, with 27 events in the four-method common set [4]. RT-PCR targets were matched to RNA-seq events before interval scoring. Structured *TRA2* targets were matched by cassette-exon and flanking splice sites after conversion to the SUPPA coordinate frame. Targets without full SUPPA structure were matched by cassette-exon coordinates, and circadian targets were matched by skipping-junction endpoints. Ties were resolved by coordinate distance, gene-name agreement, and read support. MAJIQ was matched at the cassette-exon level.

### Comparator intervals and statistics

MAJIQ v3, betAS, and rMATS-derived comparator intervals were evaluated on the same matched events. MAJIQ posterior means and standard deviations from ‘deltapsi.tsv’ were converted to symmetric Gaussian 95% intervals. betAS intervals were generated with the betAS R package from matched per-replicate rMATS count tables. The rMATS comparator was a Welch Student-t interval computed from per-replicate rMATS IncLevel values after orientation correction. Recalibration ablations used the same five-fold absolute-residual conformal procedure for each caller. Coverage was defined as the fraction of matched events whose interval contained paired validation ΔPSI measurement. Wilson 95% confidence intervals were reported for coverage, and paired coverage differences were tested with exact McNemar tests. Sharpness was summarized by mean interval width and the Gneiting-Raftery interval score. For the interval score, each event contributed its interval width plus a penalty when the paired validation measurement fell outside the interval. Scores were averaged across matched events, and lower values indicate sharper intervals with fewer or smaller coverage failures. Auxiliary detection analyses were limited to the *TRA2* positive/negative target set, where sensitivity, false-positive rate, and Matthews correlation coefficient were used to summarize binary event selection.

### Application analyses

Two public applications datasets were analyzed with the packaged calibrator. The DM1 analysis used the public GSE201255 rMATS skipped-exon table and sample-order file, covering 58 DM1 skeletal-muscle samples and 28 controls [15]. Events were analyzed with ‘braid differential’ using 500 posterior draws, minimum support 20 per group, and |ΔPSI| ≥ 0.1, then ranked by calibrated distance from zero weighted by the posterior probability of a large effect. The ESRP analysis used GSE64357, a mouse Esrp1/Esrp2 double-knockout versus wild-type comparison, to evaluate recovery of known epithelial splicing targets [16].

### Software, reproducibility, and ethics

BRAID is implemented in Python and available at https://github.com/kangk1204/BRAID. The repository includes the braid command-line package, benchmark scripts, committed result files, and the packaged differential calibrator. This study reanalyzed public RNA-seq, RT-PCR, and long-read sequencing datasets. No new human or animal specimens were collected, and no participant-level identifiable information was used.

## Results and discussion

### How BRAID builds a calibrated interval

Splicing callers can rank candidate events by statistical evidence, but this does not directly state how reliable a reported RNA-seq ΔPSI value is against an orthogonal validation assay. We designed BRAID to address this question by adding an RT-PCR-calibrated interval to an RNA-seq ΔPSI estimate (Fig 1). In the rMATS example, the input is a cassette-exon event with rMATS inclusion and skipping counts, and ΔPSI is defined as the change in percent-spliced-in between two conditions (Fig 1A). Here, ΔPSI is estimated from the rMATS inclusion and skipping counts; rMATS FDR and IncLevelDifference are retained in the output. BRAID then returns a 95% interval calibrated to RT-PCR measurements. The result is a reliability statement attached to the RNA-seq ΔPSI estimate. The interval width is learned from calibration events with paired RNA-seq and RT-PCR ΔPSI values. For each event, BRAID records the absolute gap between the two measurements. The empirical distribution of these gaps defines the half-width, *q*, used for the calibrated interval (Fig 1B). For the packaged differential-splicing calibrator, this value is *q* = 0.341, estimated from 162 RT-PCR-validated skipped-exon events from the *TRA2* and circadian calibration datasets [5,7,12,13]. A new event is then reported as the RNA-seq point estimate plus or minus this calibrated width, with bounds limited to the valid ΔPSI range. This calibration changes the interpretation of the interval, not the point estimate. A sampling-only RNA-seq interval can be narrow when within-experiment variability is small, but it may still fail to contain the RT-PCR measurement (Fig 1C). BRAID leaves the RNA-seq estimate as the center of the interval and sets the width according to the empirical RNA-seq-to-RT-PCR residual scale. In this way, the interval represents calibrated coverage against the validation assay rather than only uncertainty within the RNA-seq experiment. BRAID therefore reports the same RNA-seq ΔPSI estimate with an interval calibrated to RT-PCR residuals.

**Fig 1.**
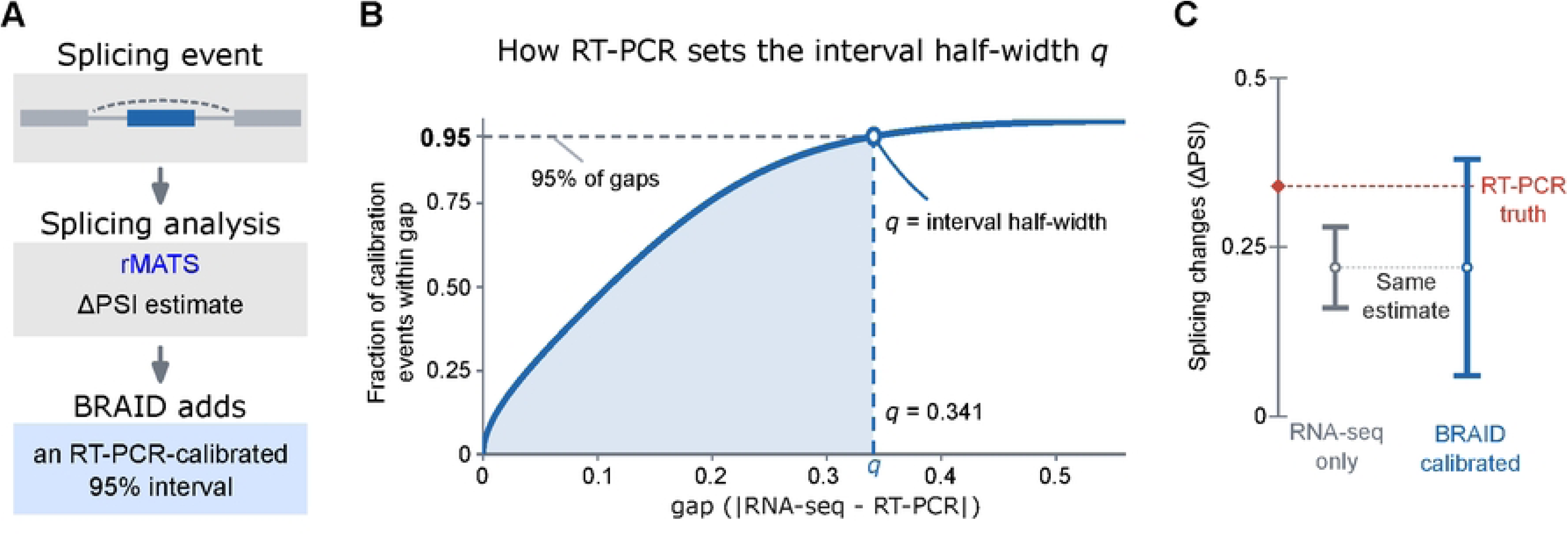
BRAID adds an RT-PCR-calibrated interval to an RNA-seq ΔPSI estimate. (A) In the rMATS workflow, a cassette-exon event is described by inclusion and skipping counts. ΔPSI is the change in percent-spliced-in (PSI) between two conditions. BRAID estimates RNA-seq ΔPSI from these counts and returns an RT-PCR-calibrated 95% interval. The panel shows the rMATS version; the same calibration can also be applied to point estimates from other splicing callers. (B) The interval half-width, *q*, is learned from calibration events with paired RNA-seq and RT-PCR ΔPSI measurements. For each event, BRAID records the absolute gap |RNA-seq − RT-PCR|. The empirical distribution of these gaps defines *q* as the 95th-percentile split-conformal residual at *α* = 0.05. A new interval is centered on the RNA-seq estimate with half-width *q*. (C) Schematic comparison of a sampling-only RNA-seq interval and a BRAID-calibrated interval on the same ΔPSI estimate. The shematic shows a calibrated interval that is wider than the sampling-only interval and contains the RT-PCR measurement. Apart from the default calibrator value, *q* = 0.341 from *n* = 162 RT-PCR-validated events, the figure is schematic.

### BRAID restored nominal RT-PCR coverage across three validation datasets

We first asked whether the calibrated interval defined in Fig 1 could restore nominal coverage of RT-PCR ΔPSI measurements across independent validation datasets. The benchmark used cassette-exon events from three validation datasets: *TRA2* knockdown in human MDA-MB-231 cells [7,12], mouse cerebellum versus liver from the circadian atlas [5,7,13], and the PC3E versus GS689 prostate EMT comparison (SRS354082) [4]. For each dataset, RNA-seq events were matched to curated RT-PCR ΔPSI measurements, oriented to the same contrast, and evaluated on the common event set available to all methods. We compared BRAID with three caller-derived interval estimates: Gaussian 95% intervals reconstructed from MAJIQ v3 posterior ΔPSI summaries [6], betAS intervals [8], and rMATS-derived Welch t intervals computed from per-replicate IncLevel values [4]. Coverage was measured as the fraction of nominal 95% intervals containing the paired RT-PCR ΔPSI value, and interval score summarized both interval width and missed coverage. Across the pooled common set of 139 events, BRAID reached 0.971 coverage, with a Wilson 95% confidence interval of 0.928 to 0.989 (Fig 2A). This was the only method whose pooled confidence interval included the nominal 0.95 target. MAJIQ, betAS, and rMATS reached 0.518, 0.734, and 0.633 coverage, respectively. At the dataset level, BRAID remained closest to the nominal target, whereas the comparator coverage estimates were consistently below 0.95. Because all methods were scored on the same events, the comparisons were paired. BRAID covered 33 events that betAS missed, whereas no event was covered by betAS and missed by BRAID in the pooled common set (exact McNemar test, P = 2.33 × 10^−10^). The same direction held against MAJIQ and rMATS, with 63 and 48 BRAID-only covered events, compared with 0 and 1 reverse-discordant events, respectively. These paired differences show that the coverage gain was not caused by a change in the evaluated event set. The higher coverage was not obtained by using an uninformatively wide interval. BRAID had the lowest pooled interval score, 0.720, compared with 2.040 for MAJIQ, 1.414 for betAS, and 1.625 for rMATS (Fig 2B). Thus, residual calibration restored RT-PCR coverage while retaining the best overall interval score among the tested interval estimates.

**Fig 2.**
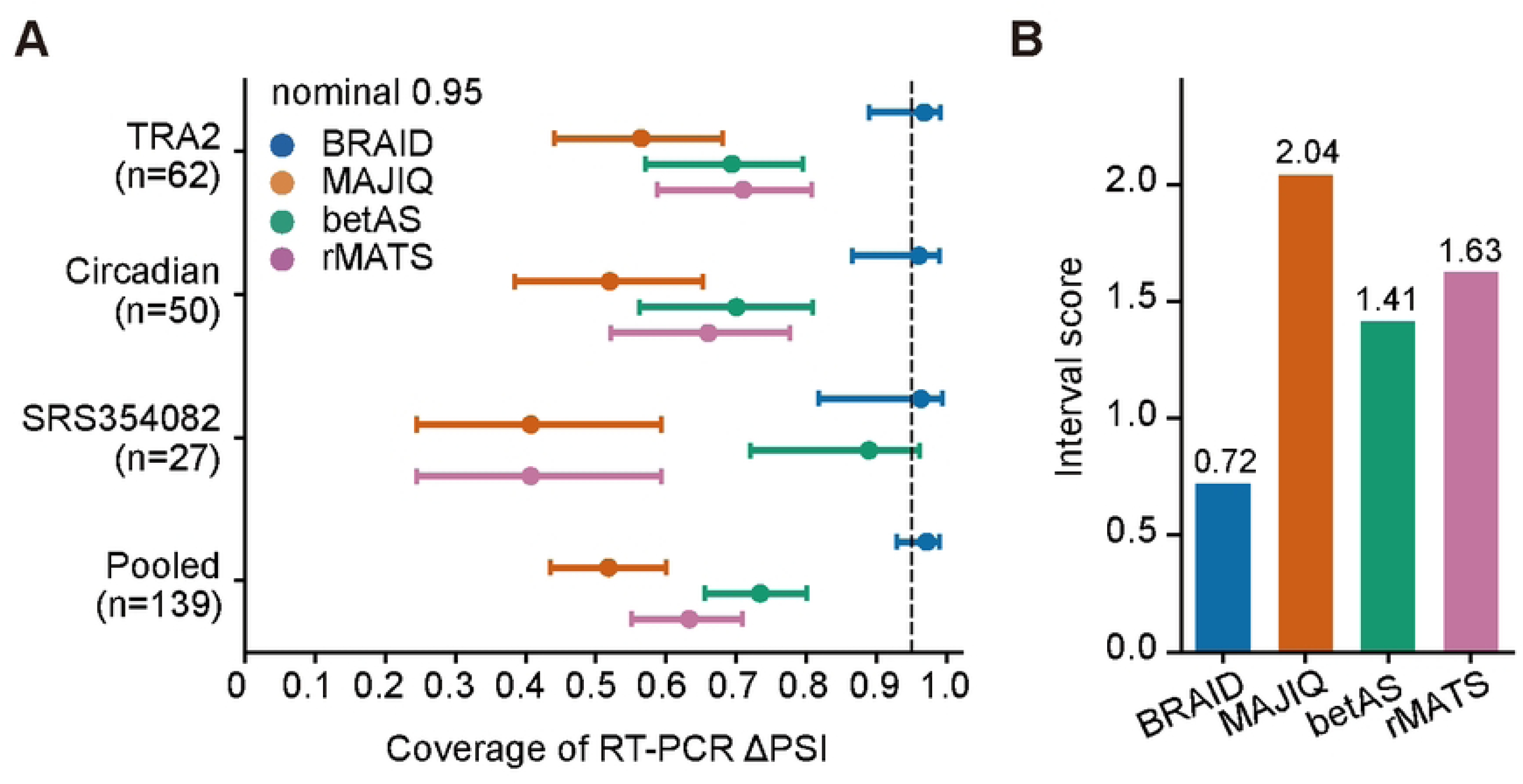
RT-PCR benchmark of calibrated interval coverage and interval score. (A) Empirical coverage of nominal 95% ΔPSI intervals on three RT-PCR validation datasets and on the pooled common set. Coverage was defined as the fraction of matched events whose interval contained the paired RT-PCR ΔPSI measurement. Points show coverage estimates and horizontal bars show Wilson 95% confidence intervals. The dashed vertical line marks the nominal 0.95 target. BRAID intervals were compared with Gaussian intervals from MAJIQ posterior summaries, betAS intervals, and rMATS-derived Welch Student-t intervals on the same matched events. Colors indicate method: BRAID, blue; MAJIQ, orange; betAS, green; rMATS, magenta. (B) Mean Gneiting-Raftery interval score on the pooled common set. Lower scores indicate sharper intervals after penalizing non-coverage. In this benchmark, BRAID had the highest pooled coverage and the lowest interval score among the compared methods.

### BRAID filters a public DM1 rMATS table to interval-supported disease events

We applied BRAID to the public GSE201255 skeletal-muscle rMATS skipped-exon table, a DM1 case-control analysis with 58 DM1 samples, including 36 congenital and 22 adult-onset cases, and 28 controls, including 21 pediatric and 7 adult controls [15]. The contrast was disease minus control. After the minimum-support filter, BRAID processed 144,975 skipped-exon events. rMATS identified 967 events with FDR < 0.05 and |ΔPSI| ≥ 0.1. Requiring the calibrated 95% interval to exclude zero reduced this set to 68 high-confidence events (Fig 3A). The high-confidence events included genes with prior evidence for DM1 or muscle-splicing dysregulation. *CLCN1* missplicing reduces ClC-1 chloride-channel function and contributes to myotonia [17,18]. *INSR* missplicing has been associated with insulin resistance in DM1 [19]. *BIN1* missplicing has been linked to T-tubule defects and muscle weakness [20], and *LDB3* splicing abnormalities have been reported in DM1 skeletal muscle with altered PKC-binding affinity [21]. The same filtered set also contained *MBNL1*, *MBNL2*, *NFIX*, *TTN*, *TNNT3*, and *NEB*, genes reported in MBNL-dependent, DM-associated, or muscle-splicing programs [21–26]. Among the displayed high-confidence events, *TTN*, *TNNT3*, *NEB*, *BIN1*, *LDB3*, *MBNL1*, *CLCN1*, *NFIX*, and *MBNL2* had positive disease-control ΔPSI intervals, whereas *INSR* had a negative interval (Fig 3B). All displayed intervals excluded zero after calibration. Thus, BRAID narrowed the public DM1 rMATS table to interval-supported events while preserving known DM1 and muscle-splicing signals.

**Fig 3.**
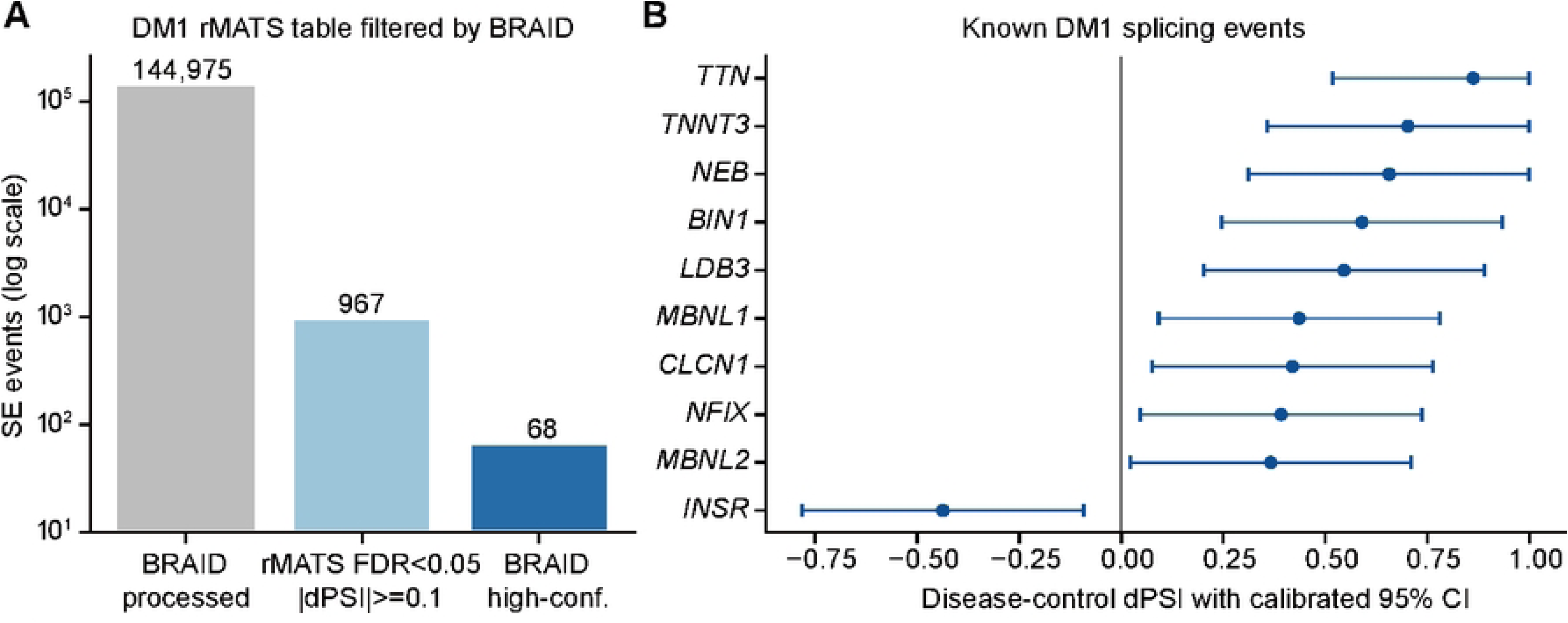
BRAID filters a public DM1 rMATS table to interval-supported events. (A) Filtering of the public GSE201255 skipped-exon rMATS table. BRAID processed 144,975 skipped-exon events. Of these, 967 passed rMATS FDR < 0.05 and |ΔPSI| ≥ 0.1, and 68 were retained as BRAID high-confidence events. (B) Recovered DM1 and muscle-splicing anchor events among the high-confidence calls. Points show disease-control ΔPSI estimates, and horizontal bars show BRAID calibrated 95% intervals. The vertical line marks ΔPSI = 0. Displayed events are high-confidence, rMATS-significant anchor events whose calibrated intervals exclude zero.

### Calibration, not stricter filtering, accounts for the coverage gain

We next tested whether residual calibration depended on the rMATS implementation used in Fig 2. For MAJIQ, betAS, and rMATS, we used the same three-dataset common set. For SUPPA2, which reports ΔPSI values but not native event-level 95% intervals, we used the *TRA2* benchmark table separately. In each case, the caller’s ΔPSI estimate was kept as the interval center, and only the interval width was recalibrated from absolute RT-PCR residuals. Recalibration brought all tested caller outputs close to nominal RT-PCR coverage (Fig 4A). On the three-dataset common set, recalibrated coverage was 0.950 for MAJIQ, 0.957 for betAS, and 0.971 for rMATS. SUPPA2 reached 0.970 in the *TRA2* benchmark after the same procedure. Thus, the coverage gain was not specific to one caller or to a favorable rMATS point estimate. The same residual calibration corrected under-coverage across different ΔPSI outputs. The gain in coverage came with a measurable increase in interval width (Fig 4B). MAJIQ, betAS, and rMATS native intervals were narrower, but all under-covered RT-PCR ΔPSI. After recalibration, mean interval widths were 0.591, 0.649, and 0.626, respectively. In the separate SUPPA2 *TRA2* benchmark, the recalibrated mean interval width was 0.691. This result separates ranking from calibration. A narrow RNA-seq-derived interval may still be useful for ranking events, but RT-PCR coverage requires an interval width matched to the empirical RNA-seq-to-RT-PCR residual. We also tested whether the packaged calibration value could be used as a hard rMATS effect-size cutoff rather than as an interval half-width. In the *TRA2* RT-PCR validation set, strict rMATS calls defined by FDR < 0.05 and |ΔPSI| ≥ 0.341 selected 17 events, with recall falling to 0.211 and MCC to 0.238 (Table 1). By comparison, BRAID effect-supported calls retained similar recall to the standard rMATS rule while giving the highest MCC in this comparison. On the same 17 strict rMATS calls, the rMATS native interval covered 12 of 17 RT-PCR measurements, whereas the BRAID calibrated interval covered 16 of 17. Thus, *q* is effective as an interval half-width, not as a hard cutoff. Together, these results show that BRAID improves RT-PCR coverage by calibrating interval width rather than by applying a stricter ΔPSI cutoff.

**Fig 4.**
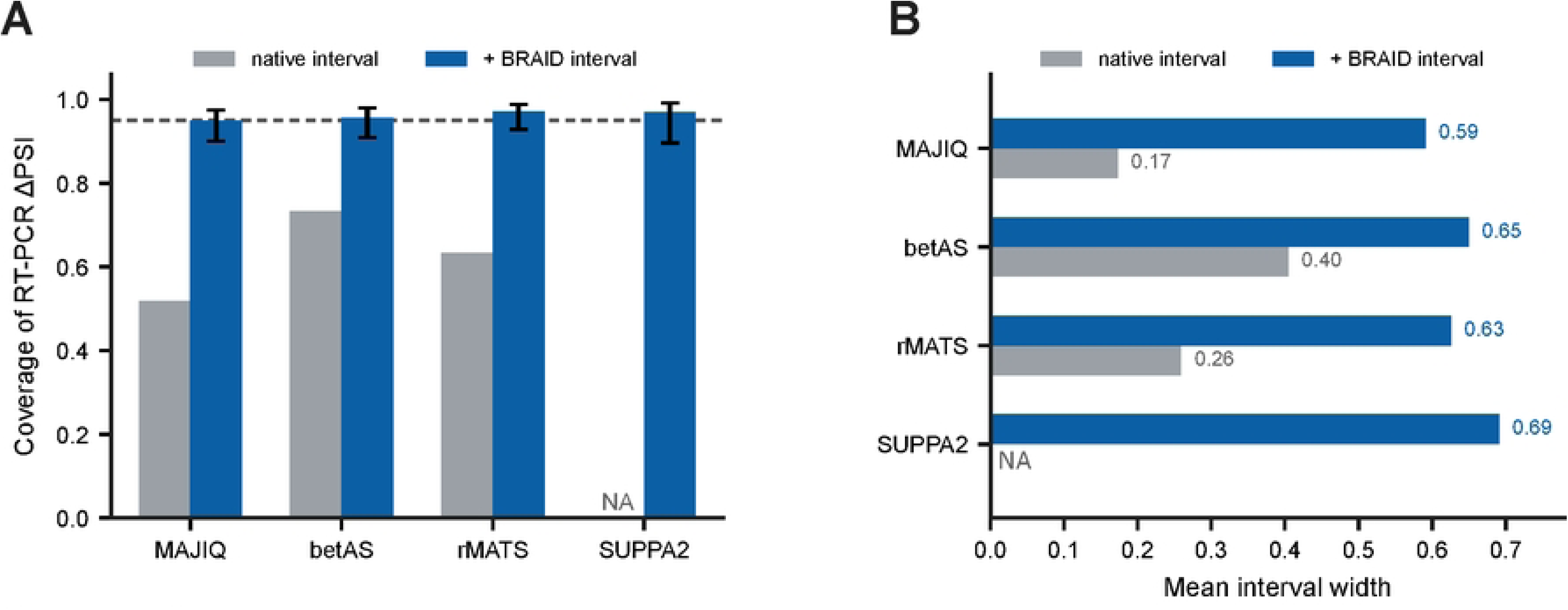
Residual calibration across splicing caller outputs. (A) RT-PCR ΔPSI coverage for native caller intervals and the same caller ΔPSI estimates after BRAID residual calibration. MAJIQ, betAS, and rMATS native intervals are shown in gray. SUPPA2 reports ΔPSI values but not native event-level 95% intervals in this comparison, so only the recalibrated interval is shown. Error bars show Wilson 95% confidence intervals for recalibrated coverage. The dashed line marks nominal 0.95 coverage. (B) Mean interval width before and after recalibration. Values printed in panel B are rounded to two decimal places.

**Table 1.**
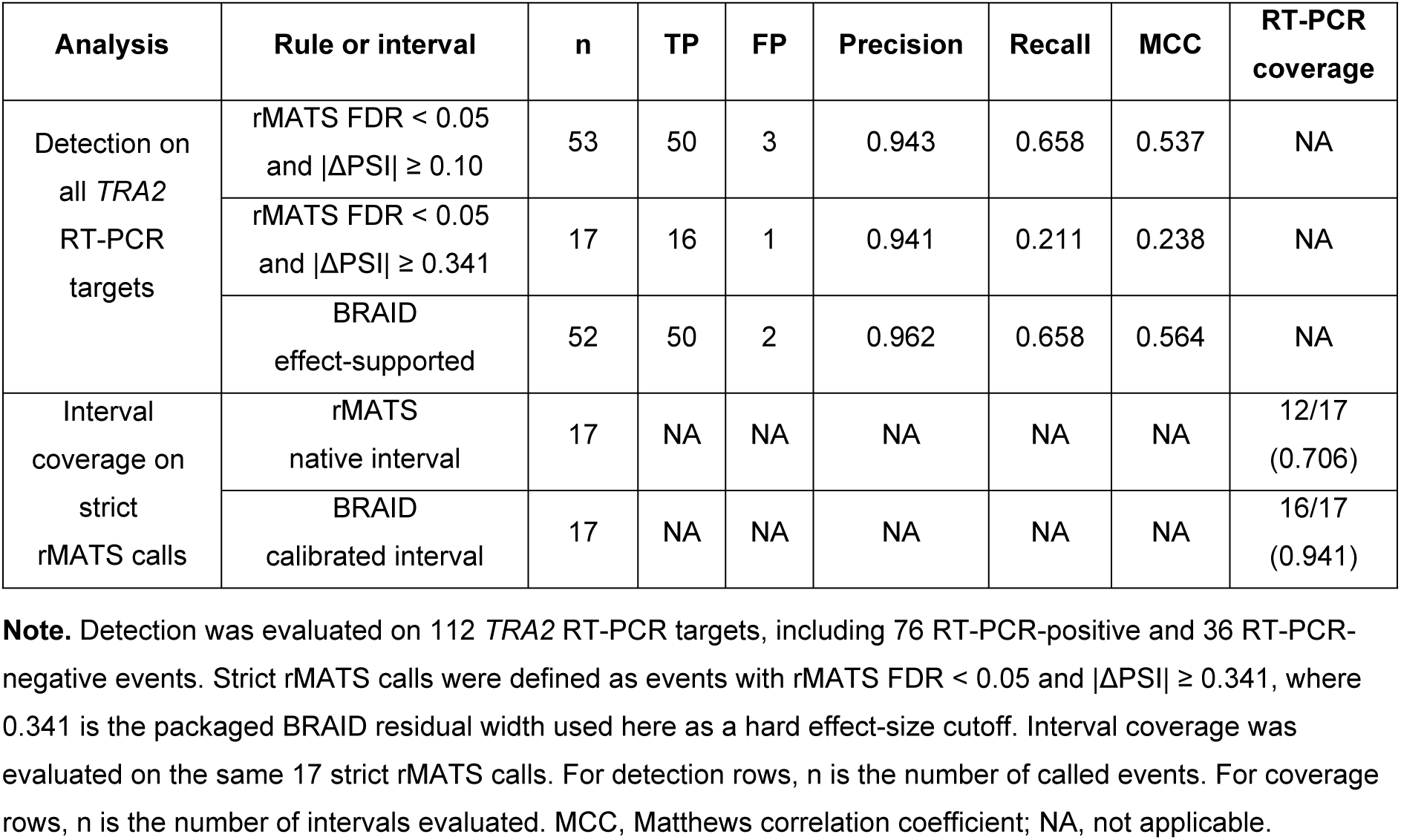
*TRA2* RT-PCR validation of strict rMATS filtering and BRAID interval calibration.

## Conclusions

BRAID adds RT-PCR-calibrated intervals to RNA-seq ΔPSI estimates. Rather than replacing an upstream splicing caller, it keeps the caller-derived event evidence and learns an interval width from empirical RNA-seq-to-RT-PCR residuals. Across RT-PCR validation datasets, this residual calibration restored nominal 95% coverage where native RNA-seq-derived intervals under-covered, while retaining the lowest interval score among the tested interval outputs. The same calibration could be applied to ΔPSI estimates from multiple splicing callers, indicating that the main gain comes from calibrating interval width rather than from a caller-specific point estimate. A stricter ΔPSI cutoff did not provide the same behavior. It reduced the call set and lost recall, whereas using the calibrated residual as an interval half-width improved RT-PCR coverage. In a public DM1 skeletal-muscle rMATS table, BRAID reduced a large set of significant splicing events to interval-supported disease events and retained known DM1 and muscle-splicing signals. BRAID is most directly useful when follow-up decisions depend on expected agreement between RNA-seq ΔPSI and orthogonal validation assays.

## Acknowledgements

This work was supported by the National Research Foundation of Korea (NRF) grant funded by the Korea government (MSIT) (NRF-2022R1A2C1093041). This work was also supported by Korea Environmental Industry & Technology Institute (KEITI) through “Core Technology Development Project for Environmental Diseases Prevention and Management,” funded by Korea Ministry of Climate, Energy and Environment (MCEE) (RS-2022-KE002173). During manuscript preparation, the authors used OpenAI ChatGPT 5.5 for limited language editing. All scientific content, analyses, and final wording were reviewed and approved by the authors, who take full responsibility for the manuscript.

## Data availability statement

BRAID source code and input-format templates are available at https://github.com/kangk1204/BRAID under the MIT license. No new sequencing data were generated in this study.

